# Understanding the infection mechanism of Φ8 using infection method in *Pseudomonas syringae* pv*. Phaseolicola* strain: *In-situ* CryoET analysis

**DOI:** 10.1101/2025.09.29.678632

**Authors:** Sanjay Kumar, Charles F. Robinson, Daija Bobe, Reza Khayat

## Abstract

The Cystoviridae family strain Φ8 is an ideal model for studying the virus-host interactions and viral assembly mechanisms. In this study, we explored Φ8 infection kinetics in P. syringae pv. phaseolicola host cells using a combination of host lysis assays, transmission electron microscopy (TEM), cryo-focused ion beam (Cryo-FIB) milling, and in-situ cryo-electron tomography (Cryo-ET). Purified Φ8 was infected with host cells and observed at different time points. The host-pathogen infection was detected using TEM studies. The infected sample underwent Cryo-FIB milling. Our findings indicated that the infected host cell membrane changes its shape during infection. We were able to visualize the various stages of Φ8 assembly using subtomogram averaging. These stages included the unpackaged procapsid (PC) having a size of ∼41 nm and the double-stranded RNA (dsRNA)-packaged procapsid having a size of ∼54 nm. The packaged PC represented the most prevalent intermediate stages during viral assembly. Notably, the majority of complete virions isolated from the host cell were found outside the host cell, revealing the extracellular predominance of fully constructed particles. These findings emphasize the importance of stable intermediate stages in viral assembly and maturation of Φ8. This work advances our understanding of viral assembly and complex host-cell interactions.

**Highlights:**

- Successfully isolated and infected *Pseudomonas syringae* pv. *phaseolicola* (*Pph*) host cells with Φ8.
- Lysis assay performed to quantify host lysis to evaluate viral infection efficiency.
- Cryo-focused ion beam milling enabled precise preparation of cellular cross-sections for structural analysis. This allows identification of key stages of the Φ8 life cycle during the infection cycle.
- The majority of complete virions were localized extracellularly, emphasizing the role of extracellular stages in the Φ8 lifecycle.

**Graphical Abstract:** 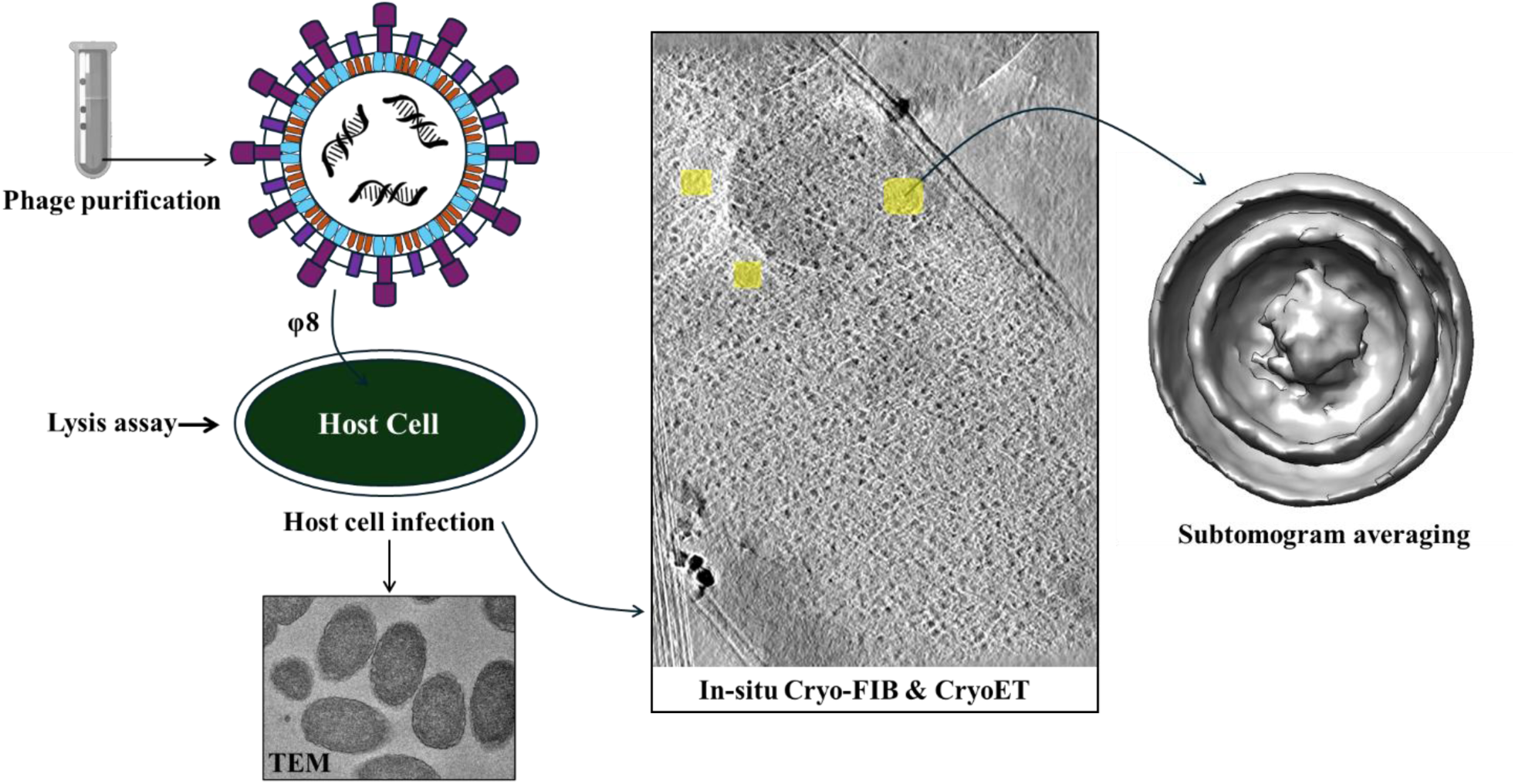

## Introduction

*Cystoviridae* represent a unique family of double-stranded RNA bacteriophages, such as Φ8, that play a critical role in shaping microbial communities by specifically targeting and infecting bacteria across diverse natural environments (Leonard et al., 1999). The *Cystoviridae* family host *Pseudomonas syringae* pv. *Phaseolicola (Pph)* is a plant pathogen (Sistrom et al., 2015). While *Cystoviridae* themselves do not infect eukaryotic organisms, their interaction with bacterial hosts can have indirect effects on human health by influencing bacterial populations and their pathogenic potential (Poranen et al., 2017). The *Cystoviridae* family is double-stranded RNA (dsRNA) virus that infect Gram-negative bacteria (Leonard et al., 1999; Mindich, 2004; Poranen et al., 2017). This family currently has seven known members (Φ6, Φ12, Φ13, Φ2954, Φ8, ΦNN, and ΦYY) (Gottlieb et al., 2002; Leonard et al., 1999). One of the most intriguing elements of the *Cystoviridae* family is its potential to undergo genetic reassortment. Because the viral genome is segmented, two different *Cystoviridae* strains can infect a single bacterial host and exchange genome segments. This reassortment can produce progeny viruses with unique combinations of genetic material, which increases their evolutionary flexibility (Mäntynen et al., 2021; O’Keefe et al., 2010; Silander et al., 2005). These viruses have unique structures, lifespan, and host interaction. Their genome is divided into three dsRNA segments known as large (L), medium (M), and small (S). The L segment contains genes for the viral RNA-dependent RNA polymerase and packaging enzymes. The M segment encodes proteins involved in the virus’s envelope and membrane fusion, whereas the S segment contains genes for capsid proteins and the entry mechanism (Jäälinoja et al., 2007). This unique feature makes *Cystoviridae* a model system for investigating viral replication (Butcher et al., 2001; Hongyan et al., 2001), assembly (Gottlieb et al., 1990), and host interactions in dsRNA viruses (Bothner et al., 1998; Doryen et al., 2005). The *Cystoviridae* Φ6 and Φ12 strains were well studied on the structure level compared with others. There is fewer structure level information available for Φ8. On the basis of similarity with other strains, Φ8 is also organized into several distinct layers. The procapsid is the innermost structure layer and houses the viral RNA. The procapsid has an icosahedral structure and is made up of various structural proteins, notably P1 (the main procapsid protein) (Mindich, 2004). The nucleocapsid shell in strains Φ6 and Φ12 encapsulates the procapsid (Huiskonen et al., 2006). However, Φ8 lacks a nucleocapsid shell(Hoogstraten et al., 2000). A bacterial host membrane-derived lipid envelope surrounds the nucleocapsid for Φ6 and Φ12 or the procapsid for Φ8 (Etten et al., 1976; Laurinavičius et al., 2004). This envelope contains viral proteins that are suggested to be required for host recognition and membrane fusion (Etten et al., 1976; Stitt & Mindich, 1983). In Φ6, the protein P8 is responsible for forming the nucleocapsid surrounding the procapsid. In Φ8, P8 is associated with the membrane envelope which surrounding the nucleocapsid and contains proteins P6, P9, and P10, which are involved in membrane fusion, envelope creation, and host cell entrance, respectively (Jäälinoja et al., 2007).

The Kainov group has previously investigated the in-vitro study of the viral assembly of Φ8 (Kainov et al., 2003). The life cycle begins with the virus binding and fusing with the bacterial outer membrane. This is facilitated by the spike complex proteins that recognize specific receptors on the host cell surface (Bamford et al., 1976; Romantschuk & Bamford, 1985). Upon binding, the virus undergoes fusion, in which the viral envelope merges with the bacterial outer membrane, allowing the nucleocapsid to enter the bacterium’s periplasmic region (Bamford et al., 1987; Caldentey & Bamford, 1992; Romantschuk et al., 1988). Once within the host, the nucleocapsid translocates through the inner membrane into the cytoplasm. While in the cytoplasm of the host, the viral RNA-dependent RNA polymerase (encoded by the L segment) starts transcription of the viral genome (Kakitani et al., 1980). This process is tightly regulated and temporally controlled, with early transcription creating mRNA for the synthesis of viral proteins required for replication, and later transcription producing genomic RNA for packaging into new viral particles (Romantschuk et al., 1988).

Furthermore, the *Cystoviridae* family’s distinct host entry features could potentially be utilized for the development of novel treatment techniques, such as phage therapy (Chambers et al., 2024; Serwer et al., 2021). Given the development in antibiotic-resistant bacterial strains, bacteriophages such as the *Cystoviridae* family are being studied as potential drug delivery systems. Phage therapy uses bacteriophages to selectively target and kill pathogenic bacteria, providing a highly precise and potentially successful treatment for bacterial illnesses.

In this study, we explore the viral assembly inside the host cell and their more stable stages. Utilizing the method of *in-situ* Cryo-ET, we can observe different life cycle stages of Φ8 inside the host. This approach helps us to explore more about the mechanism of model organism assembly, structural features and utilization as a potential drug delivery system.

## Methodology

### Phage purification

*Pph* (ATCC #21781) frozen stocks (80 % (v/v) glycerol) were plated on Luria-Bertani (LB) plates for 36 hours at 25°C. A colony was then placed in 1 L of LB broth and grown to an OD_600_ of 1.2. The stationary phase culture was then centrifuged at 5,000 x g for 15 minutes at 4°C. The resulting pellet was then resuspended in 50 ml of LB broth. Φ8 at a PFU of 2.7 x 10^11^ PFU ml^-1^ was incubated with the concentrated host for 15 minutes. The phage-host mixture was then incubated with top agar (0.8% (w/v) agar) and plated on LB plates. The plates were then incubated for 16 hours at room temperature. The top agar was then scraped from the plates and centrifuged at 26,000 x g for one hour at 4°C. The supernatant was then removed and centrifuged at 26,000 x g for one hour at 4°C. The supernatant was then ultracentrifuged at 100,000 x g for 1 hour at 4°C. The pellet was resuspended in P buffer (10 mM potassium phosphate, 1mM Mg_2_Cl, pH 8.0). The resuspended pellet was then placed on a sucrose gradient of 10 % (w/v) – 30% (w/v) and centrifuged at 63,400 x g at 1 hour (Beckman Coulter SW 40 Ti). The fraction containing the phage was then placed on a sucrose gradient of 40 % (w/v)-60 % (w/v) and centrifuged at 63,400 x g at 1 hour (Beckman Coulter SW 40 Ti). Dialysis was performed to remove the sucrose from the phage sample. The 1 ml fraction containing the phage was placed in a 50 kDa dialysis membrane (Spectrum Labs). The dialysis tube was then placed in a 500 ml buffer of 20 mM KH_2_PO_4_, 50 mM NaCl, pH 7.5 for 16 hours at a temperature of 4°C. The phage sample was placed in a 1 kDa dialysis membrane (Spectrum Labs) to concentrate on the sample. The dialysis membrane containing the sample was placed in a 500 ml buffer of 10% (w/v) polyethylene glycol 3350, 20 mM KH_2_PO_4_, 59 mM NaCl, pH 7.5 for 12 hours at a temperature of 4°C. The purified phage was then stored in -80°C in 80% (v/v) glycerol.

### Lysis Assay

A 50 ml culture of *Pph* grew to an OD_600_ of 1.5 at 25°C. A 3 ml culture of the stationary phase host was then infected with Φ8 at a multiplicity of infection (MOI) of 15. The absorbance of the phage-host mixture at an optical density of 600 was then reported every 70 minutes using a VWR® UV-1600PC Spectrophotometer.

### Transmission Electron Microscopy

*Pph* cells were infected with Φ8 at a MOI of 15 for 420 minutes. The phage-host mixture was centrifuged at 5,000 x g for 15 minutes. The pellet was then resuspended and fixed in 4 % (v/v) glutaraldehyde, 2 % (v/v) paraformaldehyde in 0.1 M sodium cacodylate, pH 7.4 overnight. The fixative was removed by centrifuging 5,000 x g for 15 minutes. The pellet was resuspended and rinsed with 0.1 M sodium cacodylate, pH 7.4. The rinsing step was repeated three times. The resulting pellet was then resuspended and fixed with 2% (v/v) osmium tetroxide in 0.1 M sodium cacodylate for one hour at room temperature. The fixed sample was then centrifuged at 5,000 x g for 15 minutes. The pellet was then resuspended and rinsed with distilled water five times. The resuspended pellet was then incubated with 1% (v/v) uranyl acetate for 1 hour at room temperature. After centrifuging the pellet was then resuspended in 30 % ethanol and dehydrated for 5 minutes. The dehydrated sample was then centrifuged again and resuspended in increasing concentrations of ethanol (50%, 70%, 80% and 90%). The sample was then dehydrated three times with 100% ethanol for 10 minutes. The sample was infiltrated with increasing parts of Spurr’s resin (3:10, 1:1, 7:10 in 100% ethanol) (Electron Microscopy Sciences). The sample was then infiltrated with 100% Spurr’s resin for six hours on a shaker. The sample was placed in a beem capsule with the accelerator (dimethylaminoethanol) in 100% Spurr’s resin. The embedded sample was cured in an 80°C oven for 16 hours.

The embedded sample was then sectioned with a microtome and placed on a carbon coated 400 mesh grids. The sectioned sample was then stained with 0.6 % (w/v) lead citrate and 2 % (w/v) uranyl acetate. Sections were observed with a JOEL 2100 TEM operating at 200 kV. Images were captured with a Gatan US1000XP at a 2000 x 2000 pixel resolution.

### Freezing of Φ8 at different time scale

The infected cells of *Pph* (ATCC® 21781™) by Φ8 at following time scale; 10-minute, 20-minute, 30-minute, 60 minutes loaded onto glow-discharged grids coated with C-flat Holey thick carbon grids (2.0μm hole, 1.2μm space with 400 mesh: Cat. # CFT412-50) (https://www.emsdiasum.com/).

### High Pressure Freezing

A Wohlwend Compact 01 high pressure freezer (HPF) was used to freeze infected cells onto grids using the Waffle method (Kelley et al., 2022; Klykov et al., 2022). The grids used were Quantifoil 2/2 200 mesh Cu, with an extra 20 nm of carbon sputtered atop the film. Type B planchettes were sanded and polished on the flat side. 1-Hexadecene was applied to the planchettes to help with separation. The waffle assembly was as follows: type B planchette with flat side facing up, grid with gridbar side facing up, sample applied to gridbar side, type B planchette with flat side facing down. This “sandwich” was assembled in the HPF tip, then frozen. The grids were then clipped using the ThermoFisher Scientific Cryo-FIB AutoGrids.

### Cryo-FIB-Milling: thinning method

An Aquilos 2+ dual-beam FIB/SEM equipped with Delmic Ice Shield and internal fluorescent light microscope (iFLM) was used for lamella preparation. A 2 kV accelerating voltage and a 13 pA electron beam current were used for SEM imaging, and a 30 kV accelerating voltage and a 10 pA ion beam current were used for Cryo-FIB imaging. Lamella were prepared as described in the Waffle Method protocol (Klykov et al., 2022). All lamellae were milled with a milling angle of 20°.

### Cryo-ET: imaging conditions

After lamellae were milled, the Autogrids were loaded into a cassette, ensuring that the milling direction of lamellae was perpendicular to the tilt axis for cryoET data collection. The lamellae were imaged using a Titan Krios microscope (ThermoFisher Scientific) operated at a voltage of 300 kV and equipped with a bioQuantum post-column energy filter (Gatan) and a K3 Summit direct electron detector (Gatan). Image acquisitions were controlled using Leginon (https://github.com/leginon-org/leginon) and SerialEM (Mastronarde, 2005). The tilt series (TS) were acquired at a nominal magnification of 33,000x with pixel size of 2.077Å. The target defocus was in the range of −1 to −8 μm. Micrographs with 31 frames were recorded with a 50ms per frame exposure time at each tilt angle (Table S1).

### Image processing

Five sets of data were collected at different time scales in an image format. All *in-situ* tomographic images converted into tilt series and processed first to eliminate bad tilt series using in-house scripts. Processed raw tilt series were selected for alignment and 3D tomogram reconstruction done by AreTomo v1.3 (Zheng et al., 2022). Aligned tomograms were reconstructed by applied tilt correction (-1), weighted back projection (WBP) and simultaneous algebraic reconstruction (SART) method. The reconstructed tomograms selected for enhancing contrast by denoising and missing-wedge correction followed by CTF deconvolution from deep learning method available in ISONET v0.2 (Y.-T. Liu et al., 2022). In these 20 random tomograms were selected for generating self-supervised deep learning model that was used for correcting all tomograms. The experimentally determined density maps of the Φ8 virion [EMD-1299], Φ8 polymerase complex/core [EMD-1300], and Φ6 nucleocapsid [EMD-0299], were selected as templates for reference-based particle picking using the EMAN2. Density map for membrane only and core with genome template created using map eraser tool from ChimeraX using Φ8 virion [EMD-1299] density map. Following the verification of particle morphology, and dimensions, these reference maps were used to visualize the particles within selected tomograms. This was achieved by overlaying the corresponding density maps onto the identified particle positions in clipped view. The host cells and Φ8 bi-layer membrane were confirmed by MemBrain-Seg tool using v10 pre-trained model. The segmentation of selected tomograms was done by ArtiaX (Ermel et al., 2022) module in ChimeraX v2.1(Pettersen et al., 2021) and visualization of tomograms using IMOD (Danita et al., 2022; Kremer et al., 1996).

### Sub-tomogram averaging

Selected preprocessed tomograms (aligned and denoised) were processed through Dynamo v1.1.552 (Castaño-Díez et al., 2012). Loaded all selected tomograms on the dynamo_catalogue and particles picked manually by selecting center vesicle. After that particle extracted with crop side-length of 30, 40, 50, 60, and 70. A low resolution ab-initio model generated with selected side-length and performed refinement using template and masks with same side-length. The refinement of complete phage particles is set by adjusting low-pass filters, angular sampling, and translation limits with different iterations (50, 40, 30) in round 1, 2, and 3. A C1 (default) and icosahedral (ico) symmetry used to check improvement of model symmetry. The reconstructed averaged structure model visualized to see the features of Φ8 using ChimeraX (Pettersen et al., 2021).

## Results and Discussion

In this study, we proposed a complete life cycle of Φ8 and their longest stable stage using cryogenic focused ion beam milling (cryo-FIB) method. Originally, we collected five sets of data that includes with and without infection of Φ8 in cells of *Pph* (ATCC® 21781™).

### Assessing Host Lysis Under Infection

To observe the life cycle of Φ8 with *in-situ* cryo-ET, we needed to see numerous phage particles inside of the host. The time point before host lysis should contain the most amount of Φ8 particles. Previously, researchers determined *Cystoviridae* lyse host cells at 70 minutes post initial infection (Kivelä et al., 2004). The cellular pellet required for *in-situ* cryo-ET needs to be 0.5 cm ≥. For *in-situ* cryo-ET, cells are harvested as a dense pellet from at least 0.5 mL of culture. We infected *Pph* with Φ8 at a multiplicity of infection (MOI) of 15. This increase in cell density allowed for a larger cell pellet size. Φ8 host lysis was first observed at 210 minutes after the initial infection. This lysis continues until 720 minutes post initial infection. From 720 minutes to 1080 minutes there was an increase in absorbance (Figure 1). This increase is due to the growth of Φ8 resistant host present in the culture. We determined that host infected cells should be fixed at 420 minutes post initial infection to visualize multiple Φ8 particles inside the host.

**Figure 1.**
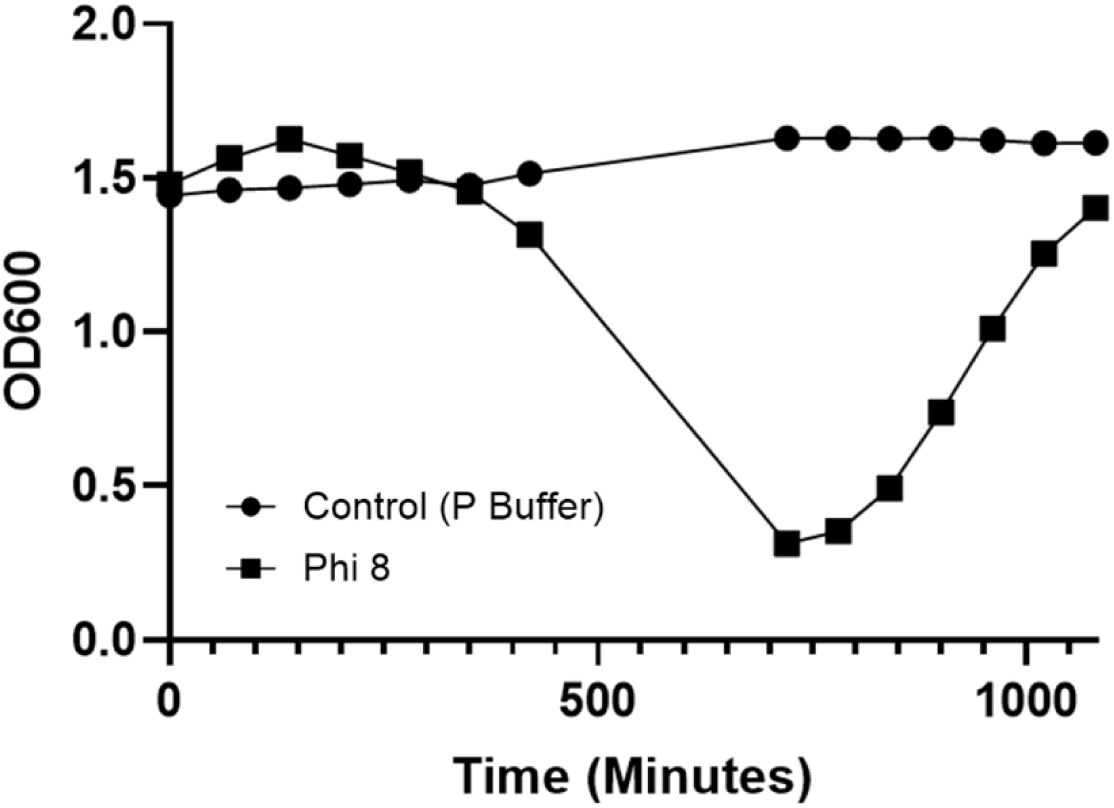
Host lysis assay with Φ8. The average OD_600_ of *Pph* infected with Φ8. n = 5. OD_600_ was monitored for ∼18 hr. Control cultures (circles) remained near OD 1.5–1.6. Φ8-infected cultures (squares) showed a ∼400–650 min lysis phase, reaching a minimum OD ∼0.25 (≈83% drop), followed by regrowth to OD ∼1.2–1.3 consistent with survival and outgrowth of uninfected cells.

### TEM Analysis of Φ8 Phage Infection in Host Cells

Thin sections of *Pph* infected by Φ8 were observed with TEM. Substantial Φ8 particles were observed intracellularly and extracellularly (Figure 2). The phage particles range from 70 nm to 96 nm (Figure 2.5). Lysate, including large vesicles, were also present in these micrographs (Figure 2.6). Large, unstained spherical aggregates were found in 70 minutes and 420 minutes post initial infection thin sections (Figure 2.3-4). No Φ8 particles were found in these unstained areas of our TEM micrographs (Figure 2.1-2).

**Figure 2.**
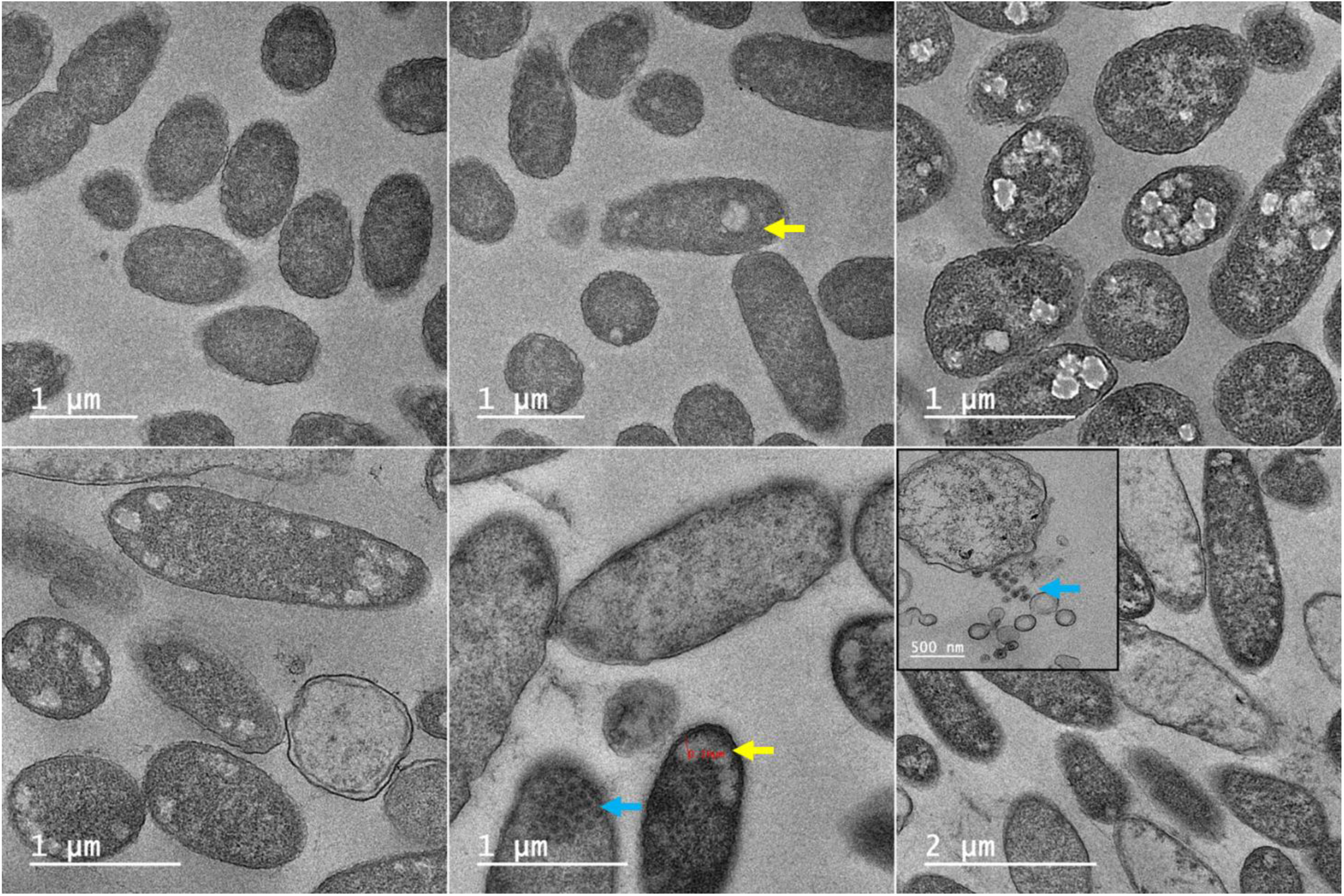
Time-resolved cryo-EM of Φ8 infection in *Pseudomonas* cells. 1. Uninfected host cells, 2. After 70hr host cells showing early infection, unstained area marked in yellow arrow, 3. Mid infection with widespread electron-dense puncta consistent with viral assembly intermediates, 4. Pre-lysis remodeling with periplasmic expansion and spheroplast-like morphology. 5 & 6. Host cells after 420 minutes showing late infection where membrane rupture (blue arrow) and ∼80–100 nm particles appeared at the cell surface; inset highlights extracellular particle clusters, and terminal lysis with disrupted cells and debris.

### *In-Situ* CryoET Analysis of Φ8 Phage Infection in Host Cells

Φ8 phage infection was investigated by infecting the host *Pph* for time intervals of 70 minutes and 420 minutes. TEM and *in-situ* CryoET experiments revealed that 70 minutes post-infection was not enough for Φ8 virion assembly within host cells. However, distinct viral assembly structures were observed at 420 minute post-infection.

### Structural analysis of Φ8

To interpret the *in-situ* structural organization of Φ8, we superimposed experimentally available cryo-EM maps of the mature Φ8 virion [EMD-1299], the Φ8 polymerase complex/core [EMD-1300], Φ6 P1P4 complex [EMD-0245], and the Φ6 nucleocapsid (EMD-0299) onto selected tomograms collected during infection. Our analysis revealed a limited number of mature Φ8 virions (∼ 3–5) within the host cytoplasm, whereas a significantly higher number (20–30) were observed outside the host cells. Superposition of the mature Φ8 particles confirmed the retention of symmetrical morphology without detectable deformation throughout the infection cycle, both inside and outside the host cells (Figure 3A). These virions exhibited a consistent, well-defined spherical shape with diameters ranging from ∼70 to 90 nm (Figure 3A). The Φ8 assembled from interactions with P4 hexamer units and a monomer unit of P1 protein. The polymerase complex (PC)/procapsid/core formed by binding P2 and P7 (De Haas et al., 1999). Each mature virion encapsulated a segmented double-stranded RNA genome within an icosahedral procapsid. Numerous procapsid particles were observed within the host cells, with occasional detection of 1– 2 procapsids outside the cell, likely resulting from the lysis or rupture of neighboring infected cells within the same tomographic field. The procapsid structure measured ∼40 nm in diameter and consisted of the P1-P4 protein complex and ∼60 additional minor structural proteins, all enclosed by a host-derived lipid envelope (Jäälinoja et al., 2007) (Figure 3A). The spatial distribution of Φ8 within the host cytoplasm suggested the presence of localized assembly zones, frequently adjacent to membrane-rich regions, which may function as scaffolding platforms or nucleation sites for virion assembly (Jäälinoja et al., 2007).

**Figure 3.**
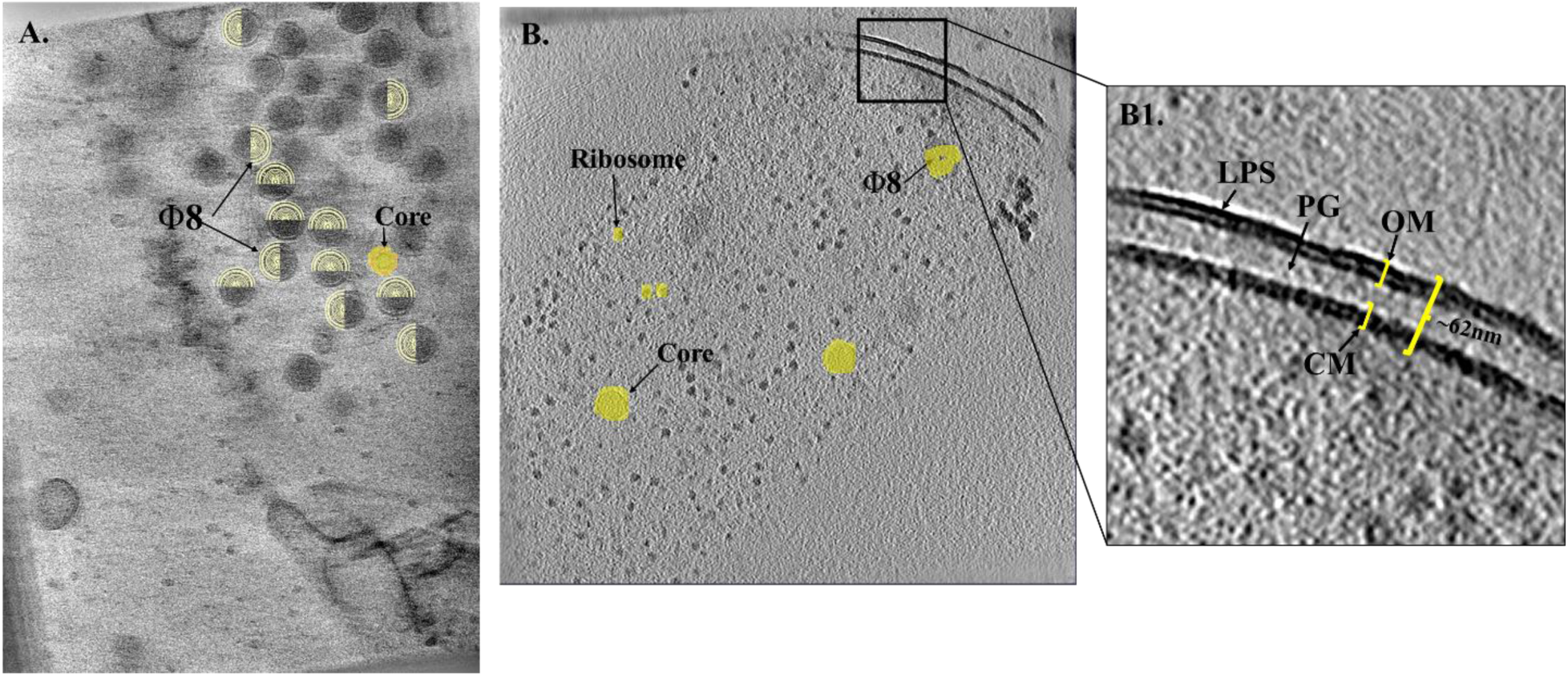
Visualization state of infected host cells. **A.** A mature φ8 particles is shown, and same virion is displayed in a clipped view, overlaid with the density map of the mature φ8 virion (yellow) and Core (magenta) for better visualization, **B.** An infected host cell, showing clearly defined membrane bilayers and visible ribosomes, **B1.** Measurements of the host cell membrane structure are highlighted: the dense outer layer is the lipopolysaccharide (LPS) region of the outer membrane which has size of ∼13 nm; the periplasmic space, which contains the peptidoglycan layer, is ∼34 nm wide; and the inner (cytoplasmic) membrane is similar in thickness to the outer membrane.

### Visual analysis of Host cell

The host cell *Pph* exhibits a consistent long, rod-shaped morphology, which remains structurally intact during both uninfected and infected states (Figure 3B). Visual analysis revealed that the host cell envelope comprises a double bilayer membrane system, consisting of an outer membrane enriched with lipopolysaccharides (LPS) measuring approximately 12 nm in thickness, an inner (cytoplasmic) membrane, and a periplasmic space containing a thin peptidoglycan layer ranging from ∼60 to 65 nm (Figure 3B1). At 70 minutes post-infection, the outer cell envelope exhibited a wave-like deformation in some cells, a feature that was less frequently observed at 420 minutes post-infection. This morphological alteration may represent an adaptive response to host immune defenses or environmental stressors aimed at mitigating viral infection. Both uninfected and infected *Pph* cells were densely populated with countable ribosomes, indicative of active cellular translation machinery. Following Φ8 infection, *in-situ* cryoET analysis allowed us to delineate the infection process into five distinct stages: (i) non-infection, (ii) initiation/adsorption, (iii) liquid-liquid phase separation (LLPS), (iv) viral assembly, and (v) host cell lysis. These stages represent the temporal and structural progression of Φ8 replication within the host cytoplasm, as visualized at nanometer resolution.

### Stage 1: Non infection stage

We observed ∼70% of host cells remained uninfected at 70 minutes post-infection, whereas only 10% at 420 minutes (Figure 4A: Stage 1). This shift suggests time-dependent increase in infection efficiency that Φ8 requires an extended period within the host to complete its intracellular replication and virion assembly processes. Based on the *in-situ* data, the majority of host cells at 70 minutes were in either the non-infected or early initiation/adsorption stages of the infection cycle (Figure 3B), with limited evidence of downstream viral activity. In contrast, the 420 minute samples showed a broader distribution of infection stages, including LLPS, viral assembly, and lysis phases. Apart from the wave-like deformations observed in the outer membrane, no major cytoplasmic changes were detected within the host cytoplasm between the 70 minute and 420 minute infection time points.

**Figure 4.**
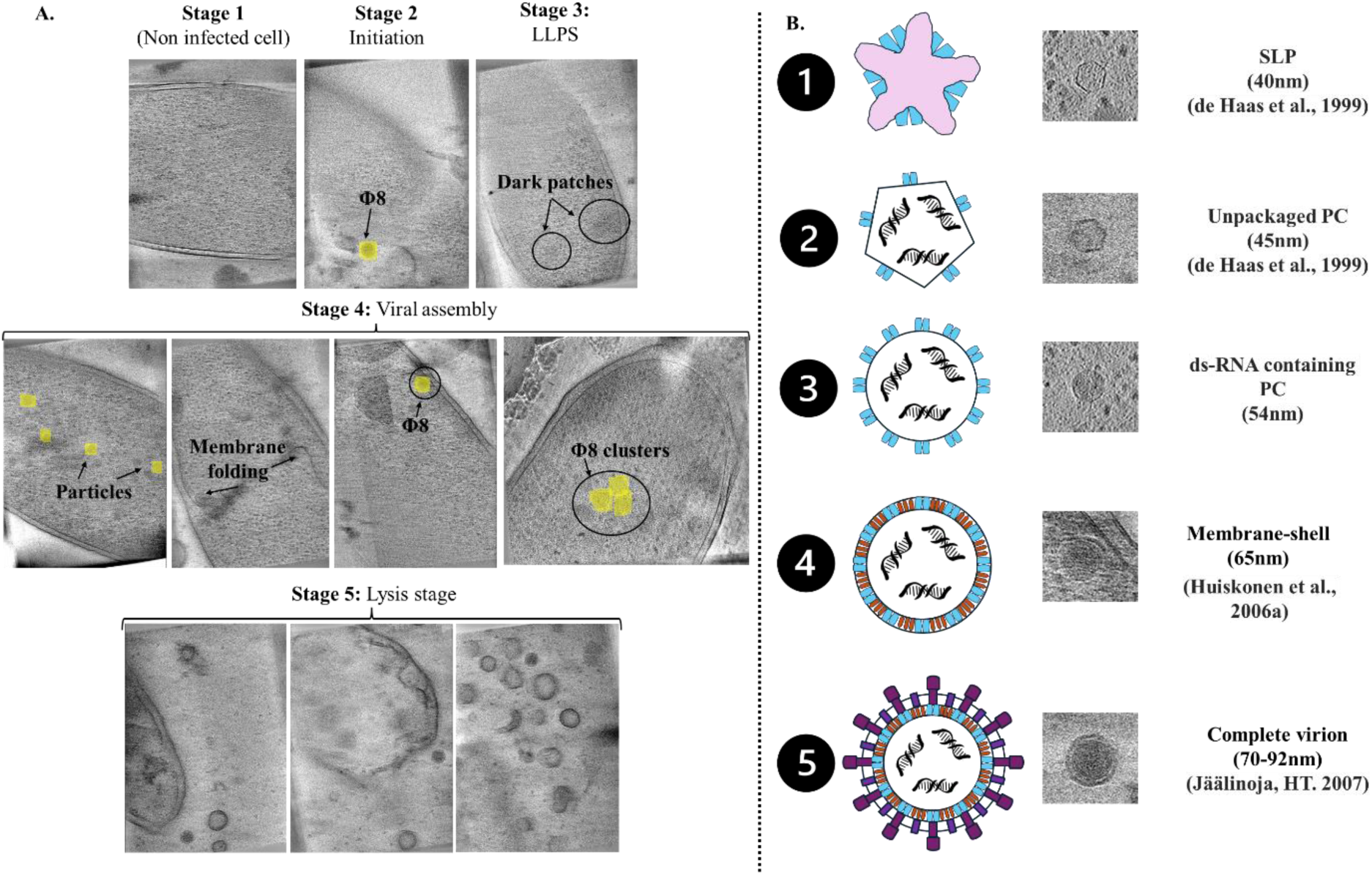
Φ8 assembly pathway in the host cell from 420 min infection cycle. **A.** Left panel: Life cycle of Φ8: **Stage 1**: Non-infected host cell. **Stage 2**: Initiation stage where Φ8 present near to host cell. **Stage 3**: LLPS stage**, Stage 4**: Viral assembly (Membrane folding and assembly happened at same time), **Stage 5**: Lysis stage where host cell first lysed then explode. **B.** Right panel: schematic representation of assembly pathway based on (Jäälinoja et al., 2007) arranged by observed intermediate stages. **1. SLP Formation:** The earliest stage observed corresponds to the formation of the S-layer particle (SLP) with an icosahedral shape. However, this stage may include structural variability, as we detected particles ranging from 41 nm to 46 nm in diameter. **2. Procapsid with dsRNA:** In the second stage, a few icosahedral particles resembling the procapsid containing double-stranded RNA (dsRNA) were observed, with an approximate size of 46 nm, consistent with earlier findings (Huiskonen et al., 2006). **3. Expanded Procapsid (PC):** The third stage involves an expansion of the procapsid, reaching a size of approximately 56 nm. This stage showed the fewest particles, suggesting it may represent a brief transition phase during intracellular maturation. **4. Membrane-Shell Assembly:** In this stage, the P8 protein assembles onto the expanded procapsid to form a membrane-associated shell, measuring around 65 nm (Huiskonen et al., 2006).We observed a small number of particles of this size located near the host cell membrane. **5. Complete Virion and Release:** The final stage likely corresponds to the mature capsid acquiring its membrane envelope, followed by host cell lysis to release fully formed virions. Only a few complete virions (∼70–72 nm) were observed inside the host cell, whereas a larger number (∼72–90 nm) were found outside, indicating active release and spread of the virus.

### Stage 2: Initiation/Adsorption Stage

The mode of host cell entry for Φ8 differs mechanistically from that of other *Cystoviridae* members, such as Φ6 and Φ12. While Φ6 typically binds to type IV pili as a primary receptor and subsequently interacts with the lipopolysaccharide (LPS) core region for membrane fusion, Φ8 appears to utilize a distinct mechanism involving direct interaction of its envelope with the truncated LPS O-antigen chains on the host outer membrane. This fusion is believed to be mediated by the viral attachment protein P3, which facilitates the delivery of the internal polymerase complex into the host cytoplasm, initiating infection (Laurinavičius et al., 2004; Mindich, 2004). We observed initiation point of host cell infection, characterized by the presence of a mature virion in close apposition to the host cell’s outer membrane (Figure 4A: Stage 2). This interaction likely represents the initiation or adsorption stage of infection. The absence of detectable host cells in the initiation phase at both 70 and 420-minute post-infection time points suggests that this stage is highly transient, likely occurring within a narrow time window that is challenging to capture using static cryo-electron tomography. This supports the hypothesis that Φ8 entry and polymerase translocation into the host cytoplasm is a rapid and efficient process.

### Stage 3: Liquid-Liquid Phase Separation (LLPS) Stage

We observed intracellular regions resembling liquid-liquid phase separation (LLPS) patterns in *Pph* host cells infected with Φ8 at both 70-minute and 420-minute post-infection time points (Figure 4A: Stage 3). These dark patches appeared as biomolecular condensates within the cytoplasm, which were not detected in uninfected *Pph* cells. While LLPS stage has not been previously reported in *Pph*, similar intracellular phase-separated condensates have been described in *E. coli* and other bacterial systems, where they are known to mediate the compartmentalization of biochemical reactions (Banani et al., 2017). LLPS enables the dynamic concentration of specific proteins and nucleic acids into membrane less compartments, often triggered by environmental fluctuations or rapid changes in cellular states (Molliex et al., 2015). In bacterial systems, LLPS has been shown to facilitate the spatial organization of RNA polymerase clusters into distinct condensates (Ladouceur et al., 2020) and plays critical roles in the assembly of membrane less organelles (Lasker et al., 2020)and in RNA metabolism (Hondele et al., 2019). Additional studies have demonstrated that LLPS is also involved in processes such as conformational rearrangements, stress response, and viral replication strategies (Boeynaems et al., 2018; X. Liu et al., 2024; Shin & Brangwynne, 2017).

In the context of Φ8 infection, the LLPS stage likely follows the initiation/adsorption phase and may serve as a prelude to viral assembly. It is plausible that the viral RNA polymerase complexes, characteristic of *Cystoviridae*, are localized and concentrated within these condensate regions to facilitate efficient transcription and genome replication. These findings suggest that Φ8 may utilize LLPS as a spatial and temporal organizer of its intracellular replication machinery.

### Stage 4: Assembly Stage

Following the LLPS stage, Φ8 proceeds to the viral assembly phase, during which viral components begin to organize into progeny virions within the host cytoplasm (Figure 4A: Stage 4). The process is initiated by fusion of the Φ8 viral envelope with the host outer membrane via lipopolysaccharide (LPS) interactions, facilitated by the viral surface protein P3 (Robinson & Khayat, 2024). This fusion event enables the release of core-associated proteins and genome segments into the host periplasmic space. Once internalized, Φ8’s RNA-dependent RNA polymerase (RdRp) P2, enclosed within the core particle, synthesizes sense strand mRNA transcripts. These transcripts associate with the host cytoplasmic membrane, which subsequently invaginates to form membrane vesicles. The viral mRNAs serve as templates for translation by host ribosomes, thus, producing essential capsid proteins, membrane proteins, and replication enzymes required for the formation of progeny particles (Poranen et al., 2017).

No evidence of viral assembly was detected at the 70-minute infection time point; however, at 420 minutes post-infection, numerous intermediate assembly states were observed within host cells. These intermediates, known as single-layered particles (SLPs) contain P1-P4 proteins to form a complex without dsRNA (Figure 5A1), represent distinct morphogenetic forms during capsid and core maturation. Although a star-shaped core morphology is common, we did not observe in our data, possibly due to its rapid transition. We identified a range of other intermediates that vary in different shapes and sizes (Figure 5A1-6). The two shapes of the polymerase core were observed; one with a hexagonal shape with dsRNA (Figure 5A2) and the other with a circular shape (Figure 5A3) which has a dense core with dsRNA and is mostly prevalent and typically arranged in clustered formations within the cytoplasm (Figure 6C1, C2A, C4A, C5A). Dense core present in high abundance suggests that this conformation may represent the most stable and long-lived state in the assembly pathway. This polymerase core is an exact match with the map of EMD-1300 and is similar to the Φ6 nucleocapsid. The core has P1 that forms a dodecahedral shell that encompasses it with P4. We observed three distinct stages of viral assembly: (i) a fully developed membrane-shell (Figure 5A4), (ii) a partially formed membrane-shell with dense polymerase core and it has a lipid bi-layer located near the host cytoplasmic inner membrane layer (Figure 5A5) and (iii) a fully assembled membrane-shell with enclosing a dense polymerase core (Figure 5.A6).

**Figure 5.**
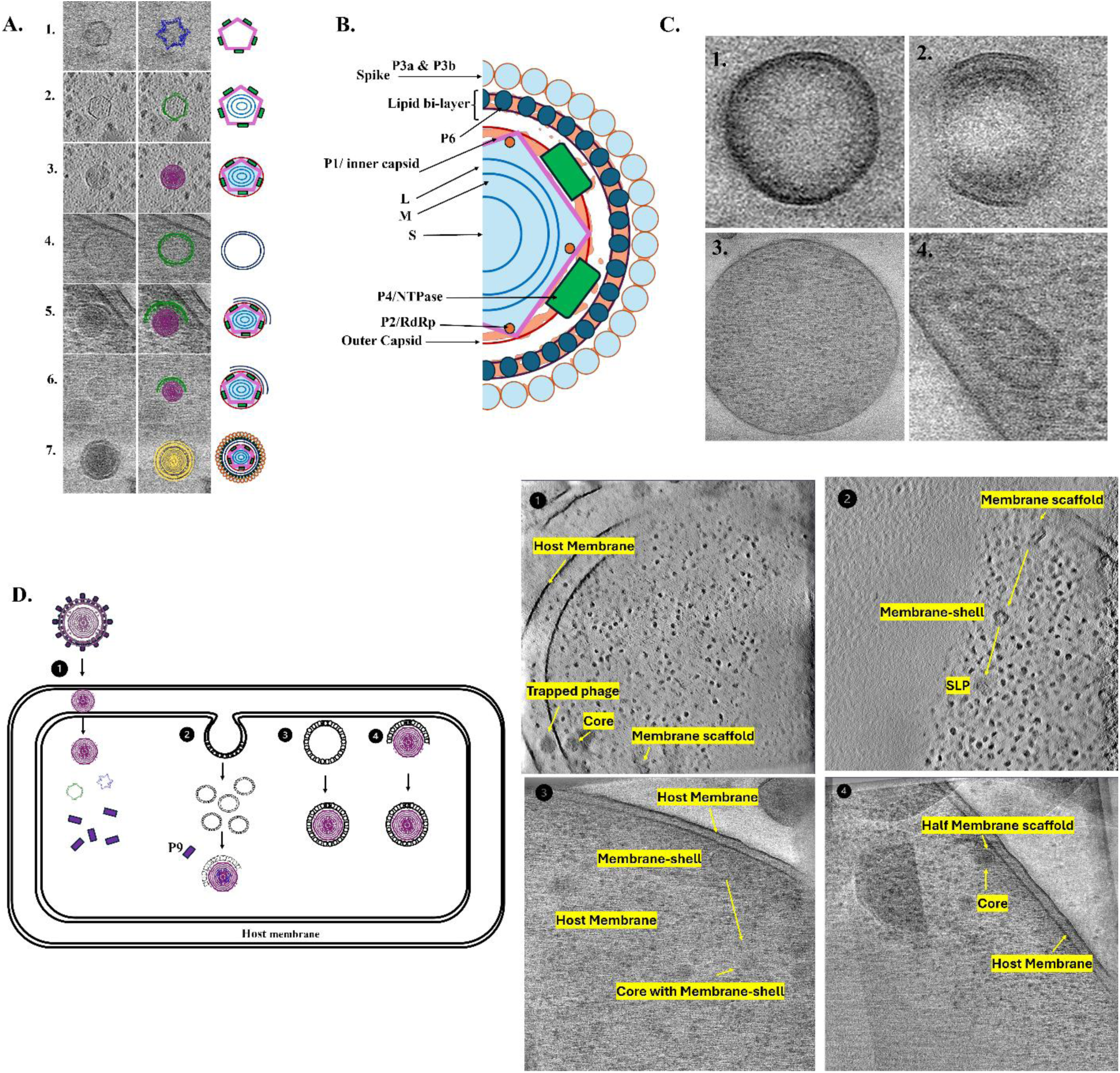
Visualization of Φ8 intermediate stages and vesicles observed during infection cycle. **A. Left panel**: **1.** Immature empty SLP with hexagonal-shaped core [template: EMD-0245], **2.** Polymerase core with genome adopted regular hexagonal shape [template: EMD-0299], **3.** Complete polymerase core adopted circular shape [template: EMD-1300], **4.** Membrane-shell present near host cell inner-membrane [template: subtract-map], **5.** Early stage of membrane-shell development that covering polymerase core [template: subtract-map], **6.** Membrane-shell maturation stage to formed vesicle [template: subtract-map], and **7.** Fully mature virions located inside host cells [template: EMD-1299]. **B**. Schematic representation of the Φ8 visible features that we observed by comparing with template map. **C. Right panel: 1**. OMV, **2.** OIMV, **3.** CMV, and **4.** IMV. D. Schematic representation of observed ways of membrane acquisition by viral particles, 1. Phage entry, 2. Process of membrane-scaffolds recruited by core particles, 3. Membrane-shell formed near host cell and later covered by core particle in the center of cytoplasm, 4. Core particle covered by membrane scaffold present near to host cell membrane.

**Figure 6.**
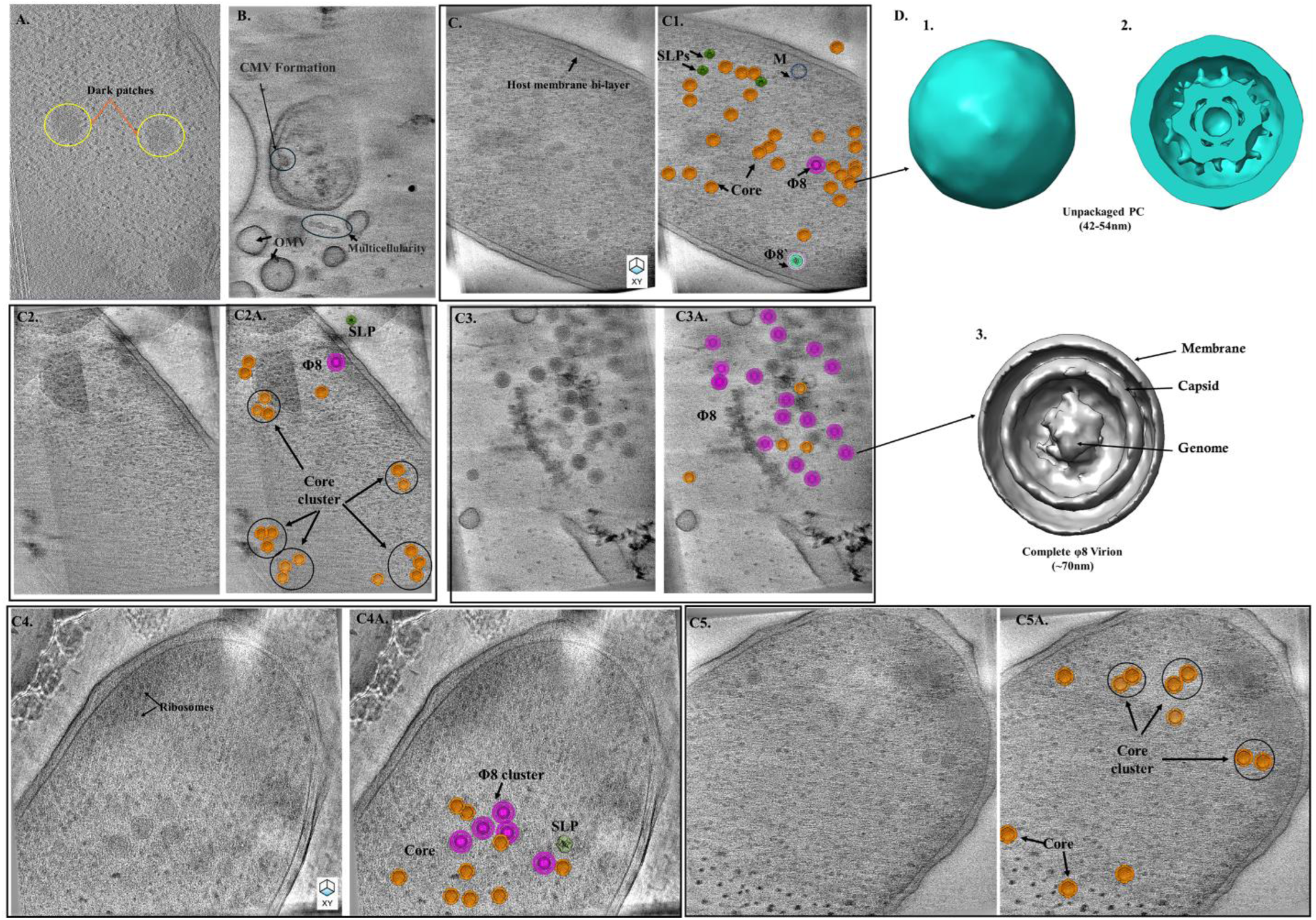
*In-situ* tomographic slice and sub-tomogram averaging analysis of Φ8 infection with host cell. **A.** liquid-liquid phase separation/ dark patches circled in yellow inside the host cell, **B.**CMV formation stage, Different sizes of OMVs, and Multicellularity pattern seen in tomogram, **C (2-5).** Φ8 assembly pathway inside the host cell. **C (C1, 2A, 3A, 4A, 5A).** Overlapping of template density maps on particle position. SLPs in green, Core in orange, Φ8 in magenta color. Cluster of particles circled circled. Sub-tomogram averaging; **1.** Initial 3D models of unpackaged PC with icosahedral symmetry, **2.** Clip of unpackaged PC, **3**. Complete Φ8 displaying membrane, capsid, and genome inside the cell.

Additionally, we observed significant cytoplasmic membrane remodeling, including ruffling and invagination, which led to the formation of cytoplasmic membrane vesicles (CMVs) and small fragments of inner membrane and inner membrane vesicles (Figure 5C1-4). The CMVs, small and large inner membrane fragments, were found dispersed throughout the cytoplasm and were often devoid of internal contents. The formation of CMVs and small membrane fragments appear to occur in parallel with viral assembly, potentially supporting virion envelopment or acting as assembly scaffolds (Figure 4A: Stage 4). Importantly, we also captured an early stage of mature Φ8 virion formation, where membrane envelopment of the core was clearly visible adjacent to the host cytoplasmic membrane, highlighting a critical transition step toward complete particle morphogenesis (Figure 5.A5). The mature Φ8 virion was observed in the center of host cell (Figure 6.A1, B1, D1). The Φ8 virion present inside the host cells have a slight ruffle in membrane while no changes were observed in those present outside the host cell or after rupturing the host cell (Figure 6C1). Following the completion of intracellular maturation, the host cells progressed to the lysis stage, culminating in the release of assembled virions into the extracellular environment.

### Stage 5: Membrane Recruitment stage

Stitt and Mindich proposed different models of membrane acquisition by Φ6 in 1983. According to their proposed mechanisms, Model I and Model II describe distinct pathways utilized by Φ6. However, our data demonstrate the presence of Model I, both subtypes of Model II, and an additional configuration resembling Model III, observed in Φ8 (Figure 5D). In Φ6, membrane acquisition is a highly coordinated and protein-mediated process. It begins with the involvement of the nonstructural protein P12, which plays a pivotal role in initiating the formation of lipid vesicles derived from the host’s inner cytoplasmic membrane. P12 facilitates the assembly of P9, a membrane-associated viral protein, into lipid-rich structures known as P9-lipid complexes (Mindich, L. et al., 1979). After the phage enters the host cell, it begins to disassemble and replicate its genome, along with synthesizing components involved in membrane recruitment (Figure 5D1). We observed that in Φ8, the viral core is fully assembled within the cytoplasm, after which it becomes associated with membrane structures. In Model I, we observed that membrane vesicles surround the viral core directly within the cytosol (Figure 5D3), forming the viral envelope without requiring proximity to the host’s inner membrane. In Model II, the viral core migrates near the host’s inner membrane, where it is wrapped by protruding lipid layers (Figure 5D4). Additionally, in Model III, membrane scaffolds appear to wrap around the core centrally within the cytoplasm (Figure 5D2).

### Stage 6: Lysis Stage

Following the maturation of Φ8 virions within the host cytoplasm, the infection cycle culminates in host cell lysis. This is characterized by the rupture of the cellular membrane and subsequent release of mature viral particles along with fragmented host membrane components into the extracellular space (Figure 4A: Stage 5). This terminal phase of infection was associated with significant structural disintegration of the host envelope, accompanied by the appearance of various membrane-derived vesicular structures and viral particles in the surrounding environment. We identified three distinct types of membrane vesicles released during or after lysis, each exhibiting unique structural features:

### 1. Outer Membrane Vesicles (OMVs)

These vesicles are composed of a single bilayer membrane, originating from the host outer membrane. OMVs typically contain lipopolysaccharides and host-derived proteins (Figure 5C1).

### 2. Outer-Inner Membrane Vesicles (OIMVs)

Structurally more complex, OIMVs consist of double bilayer membranes, encapsulating cytoplasmic content including genome fragments and occasionally mature Φ8 virions. These vesicles likely result from the combined detachment of outer and inner membranes (Figure 5C2).

### 3. Cytoplasmic Membrane Vesicles (CMVs)

CMVs are enclosed by a single bilayer membrane, derived primarily from the inner (cytoplasmic) membrane. These vesicles often contain genomic material and internal membrane fragments (Figure 5C3).

All three vesicle types varied in size and morphology, suggesting heterogeneity in their biogenesis. Based on the observed vesicle shapes and associated cellular disruption, we propose that the lysis process can be subdivided into two distinct mechanistic stages: blebbing and explosive lysis.

- In the blebbing stage, the cell membrane forms irregular bulges or protrusions, releasing vesicles in a gradual and structured manner. All three types of vesicles (OMVs, OIMVs, and CMVs) may be formed during this phase.
- The explosive lysis stage is triggered by phage-derived lytic enzymes, such as endolysins, which enzymatically degrade the peptidoglycan layer, leading to catastrophic cell envelope failure (Figure A: Stage 5). This results in the abrupt release of vesicles predominantly OMVs and CMVs which typically exhibit spherical, bubble-like morphologies and contain host cytoplasmic content and viral particles.

OMVs and CMVs were predominantly circular and varied in diameter, whereas OIMVs often appeared as multi-lamellar or nested vesicles with complex internal content, including mature Φ8 virions. The structural diversity of these vesicles suggests a role not only in viral egress but potentially in intercellular communication or immune evasion mechanisms, as previously reported in bacterial-phage systems.

### Sub-tomogram averaging

After visual inspection, we performed sub-tomogram averaging by extracting the Φ8 core and mature Φ8 virion particles (Ladouceur et al., 2020b; Jingwen & Joseph, 2020). Many Φ8 particles found in our tomograms were located near one another (Figure 3A; 4A:Stage4; 6C4, 6C4A). These particles were identified as protein capsids. The clustering of these particles could allow for rapid *Cystoviridae* transcription and assembly (Figure 6 B1, E1).

TEM micrographs and *in-situ* tomograms of Φ8 infecting *P. syringae* had a significant amount of cell lysate. These artifacts of phage infection contained many different shapes and sizes. Previously, researchers observed *P. aeruginosa* lysed cells with *in-situ* cryo-ET. Tomograms of the lysed cells revealed a consistent 35 nm crown structure (Kaplan et al., 2021). No crown structures were observed in our *in-situ* study.

## Conclusion

Our findings shed light on essential elements of *Cystoviridae* strain Φ8 infection kinetics and assembly in the host system. According to our *in-situ* findings, the unpackaged particle and dsDNA-packaged procapsid are the most stable stages of the Φ8 viral lifecycle. The prevalence of intact virions outside the host cell suggests a mechanism favoring extracellular assembly or release. This information could help to influence future research on viral assembly and infection mechanisms, potentially aiding in the development of antiviral treatments targeting comparable pathways.

## Author Contributions

**SK:** Methodology, Data curation, Validation, Visualization, Writing – original draft, review & editing. **CR:** Methodology, Data curation, Validation, Writing – original draft, review & editing. **DB:** Methodology**, RK:** Supervision, Writing – original draft, review & editing.

## Acknowledgements

We acknowledge Imaging facilities at ASRC, and City College. We also acknowledge NYSBC for data collection. We thank CCNY Microscopy Facility for access to the instrumentation needed for these studies.

---

**Table S1.** *In-situ* CryoET data collection parameters.

| **Session name** | **21sep22d** | **21oct06b** | **22mar25a** | **23sep20a** | **S23nov20a** |
| --- | --- | --- | --- | --- | --- |
| **Time of infection** | **70 minute** | **70 minute** | **70 minute** | **70 minute** | **420 minute** |
| **Microscope** | Krios 2-EF | Krios 2-EF | Krios 2-EF | Krios 2-EF | Krios 2-EF |
| **Data collection** | Leginon | Leginon | Leginon | SerialEM | SerialEM |
| **Accelerating voltage** | 300kV | 300kV | 300kV | 300kV | 300kV |
| **Total dose (e<sup>-</sup>/Å<sup>2</sup>)** | 3.71 | 4.45 | 2.78 | 3.53 | 3.45 |
| **Pixel size (Å)** | 2.077 | 2.077 | 2.077 | 2.077 | 2.077 |
| **Defocus (μm)** | -3 to -6.19 | -3 to -8.20 | -5 | -4.64 to -6.12 | -1 to -5 |
| **Magnification** | 33000 | 33000 | 33000 | 33000 | 33000 |
| **Number of tilt series** | 12 | 53 | 64 | 11 | 98 |
| **Tilt-angles** | -29 to 50 | -38 to 59 | 38 to -57 | -29 to -59 | -30 to 59 |

